# The structural and electrophysiological properties of progesterone receptor expressing neurons vary along the anterior-posterior axis of the ventromedial hypothalamus and undergo local changes across the reproductive cycle

**DOI:** 10.1101/2021.02.02.429415

**Authors:** Inês C. Dias, Nicolas Gutierrez-Castellanos, Liliana Ferreira, Susana Q. Lima

## Abstract

Sex hormone levels continuously fluctuate across the reproductive cycle, changing the activity of neuronal circuits to coordinate female behavior and reproductive capacity. The ventrolateral division of the ventromedial hypothalamus (VMHvl) contains neurons expressing receptors for sex hormones and its function is intimately linked to female sexual receptivity. However, recent findings suggest that the VMHvl is functionally heterogeneous. Here, we used whole cell recordings and intracellular labeling to characterize the electrophysiological and morphological properties of individual VMHvl neurons in naturally cycling females. We found that the properties of progesterone receptor expressing (PR+) neurons, but not PR- neurons, depended systematically on the neuron’s location along the anterior-posterior axis of the VMHvl and the phase within the reproductive cycle. Prominent amongst this, the resting membrane potential of anterior PR+ neurons decreased during the receptive phase, while the excitability of medial PR+ neurons increased during the non-receptive phase. During the receptive phase of the cycle, posterior PR+ neurons simultaneously showed an increase in dendritic complexity and a decrease in spine density. These findings reveal an extensive diversity of local rules driving structural and physiological changes in response to fluctuating levels of sex hormones, supporting the anatomical and functional subdivision of the VMHvl and its possible role in the orchestration of different aspects of female socio-sexual behavior.

## Introduction

Female reproductive physiology and behavior are under the control of the ovarian sex hormones estrogen and progesterone, whose fluctuating levels act reversibly in the female brain, organizing the activity of neural circuits to synchronize sexual behavior with reproductive capacity (Jennings & de Lecea, 2020; Snoeren, 2018). The ventrolateral division of the ventromedial hypothalamus (VMHvl) is crucial for female reproductive behavior, in particular for the display of lordosis, the female acceptance posture. Non-specific electrolytic lesions (Pfaff & Sakuma, 1979a) or ablation of genetically delineated neuronal populations (Rissman et al., 1997; Yang et al., 2013) of the VMHvl virtually abolish lordosis, while electrical stimulation at the same location enhances the probability of females displaying it (Pfaff & Sakuma, 1979b). Importantly, VMHvl neurons have rich expression of receptors for estrogen (ER) and progesterone (PR) and therefore are sensitive to the fluctuating levels of these sex hormones across the reproductive cycle (Jennings & de Lecea, 2020; Snoeren, 2018). In accordance, local infusion of sex hormones in the VMHvl increases female receptivity (Rubin & Barfield, 1983a) while male-evoked activity in this hypothalamic nucleus is enhanced when females are sexually receptive (Nomoto & Lima, 2015). Moreover, the output of PR-expressing neurons of the VMHvl is altered across the reproductive cycle and this cyclic remodeling is fundamental for the expression of acceptance when females are sexual receptive (Inoue et al., 2019). In summary, vast evidence highlights the importance of the VMHvl for coordinating female’s reproductive state with sexual receptivity.

Several studies have shown a large diversity of neuronal types with a wide variety of cellular identities within the VMHvl (Flanagan-Cato, 2011; Kim et al., 2019; McClellan et al., 2006). In addition, the generation of Cre lines providing access to ER- and PR-expressing neurons uncovered further anatomical and functional subdivisions across the nucleus’ anterior-posterior axis (AP axis) (Flanagan-Cato, 2011; Hashikawa et al., 2017; Inoue et al., 2019; Kim et al., 2019; Lo et al., 2019; McClellan et al., 2006; Wang et al., 2019), revealing a much broader role of this hypothalamic region in the control of different aspects of female socio-sexual behavior. While regulation of female sexual receptivity seems to be localized to the most posterior-lateral part of the VMHvl, its posterior-medial division is involved in aggressive behavior towards intruders in mothers (Hashikawa et al., 2017). Moreover, it was recently shown that ER-expressing neurons in the anterior portion of the VMHvl are important for self-defense in males (Wang et al., 2019). The connectivity of ER-expressing neurons also varies across the AP axis, further supporting the idea of topographic heterogeneity within this small nucleus (Lo et al., 2019).

How fluctuating levels of sex hormones affect the properties and function of VMHvl neurons remains poorly understood. Most studies aimed at elucidating the impact of sex hormones on neuronal activity in this region have focused on the structural properties of VMHvl neurons. The majority of neurons in the VMHvl have a long primary dendrite (LPD) that extends outside the nucleus, receiving inputs from other forebrain regions, a short primary (SPD), and secondary dendrites that may integrate local inputs (Calizo & Flanagan-cato, 2000; Millhouse, 1973). Previous studies in rodents have shown that externally primed estradiol treatment shortens LPDs and reduces dendritic density in the VMHvl, effects which are reversed by progesterone (Griffin et al., 2010; Griffin & Flanagan-Cato, 2008). In addition, externally applied ovarian hormones increase the density of dendritic spines on the SPD of VMHvl neurons (Calizo & Flanagan-cato, 2000), and decrease the spine density on the LPD of VMHvl neurons (Calizo & Flanagan-cato, 2002). Moreover, treatment with estrogen was found to lead to an increase in the size of neuronal somata in the VMHvl (Carrer & Aoki, 1982; Griffin & Flanagan-cato, 2011). More recently, it was shown that elevation in the concentration of circulating estrogen increases the number of axon terminals of PR-expressing VMHvl neurons that project to the anteroventral periventricular nucleus of the hypothalamus (Inoue et al., 2019). In contrast, the electrophysiological properties of VMHvl neurons across the reproductive cycle have received very little attention (but see Kow and Pfaff, 1985 and Booth and Wyman, 2010).

A noteworthy common factor among the vast majority of previous studies on the effects of sex hormones on VMHvl neurons and behavior is that they have been performed in ovariectomized females in which sex hormones were systemically administered at concentrations that do not always match the physiological levels that are observed in naturally cycling females (Liu & Shi, 2015). Moreover, despite the growing evidence suggesting anatomical and functional diversity of the VMHvl, most studies have been largely focused on its most posterior levels (Hashikawa et al., 2017; Inoue et al., 2019; Yang et al., 2013). However, the VMHvl extends slightly more than a millimeter across the base of the hypothalamus (Kim et al., 2019; Lo et al., 2019), and sex hormone receptor expressing neurons are homogeneously distributed throughout (Kim et al., 2019; Sá & Fonseca, 2017). If and how the structural and physiological properties of VMHvl neurons along its AP axis vary across the female reproductive cycle remains elusive.

In order to gain a more complete understanding of the function of the VMHvl, in the present study we investigated the structural and electrophysiological properties of genetically delineated neurons along the VMHvl AP axis and across the reproductive cycle of naturally cycling females. We focused our efforts on neurons expressing the progesterone receptor, as they are fundamental for female sexual behavior (Yang et al., 2013). To do so, we obtained slices from receptive and non-receptive females and performed in vitro whole-cell recordings of PR expressing (PR+) and PR non-expressing (PR-) neurons, with subsequent reconstruction of the recorded neurons to characterize their electrophysiological and structural properties.

Here we report a wide variety of structural and physiological properties across the AP axis of the VMHvl, which are specific for PR+ neurons and that vary locally across the reproductive cycle. For instance, the membrane resting potential of anterior PR+ neurons decreases during the receptive phase, while the excitability of medial PR+ increases when females are non-receptive. During the receptive phase of the cycle, posterior PR+ neurons simultaneously undergo an increase in dendritic complexity and a decrease in spine density. These findings reveal an extensive diversity of local rules driving structural and physiological changes in response to fluctuating levels of sex hormones, supporting the anatomical and functional subdivision of the VMHvl and its possible role in the orchestration of different aspects of female socio-sexual behavior.

## Methods

### Animals

Data was collected from adult PR-Cre-R26R-EYFP (for short PR-EYFP) female mice (2-9 months). PR-EYFP mice express the Enhanced Yellow Fluorescent Protein (EYFP) in cells expressing progesterone receptor (PR+) using the Cre-lox recombination system. Briefly, these mice result from the cross-breeding of B6129S(Cg)-Pgrtm1.1(Cre)Shah/AndJ75 mice (for short PRCre; JAX stock #017915; Yang et al., 2013), containing the Cre recombinase, which is expressed simultaneously with endogenous PR gene, with B6.129X1-Gt(ROSA26Sortm1(EYFP)Cos/J177 mice (for short R26R-EYFP; JAX stock #006148; Srinivas et al., 2001) that have the EYFP gene following a STOP sequence flanked by loxP. Therefore, in their offspring, heterozygous for both PR-Cre and R26R-EYFP alleles, loxP cassettes are removed by the action of Cre recombinase specifically in cells expressing PR, resulting in EYFP expression.

For the visualization of GABAergic somas, we used double heterozygous mice that resulted from the cross-breeding of B6J.129S6(FVB)-Slc32a1^tm2(cre)Lowl^/MwarJ (for short Vgat-ires-Cre knock-in; JAX stock #028862; Vong et al., 2011) with B6;129S6-Gt(ROSA)26Sor^tm9(CAG-tdTomato)Hze^/J (for short Ai9; JAX stock #007905; Madisen et al., 2010). Using the previously described Cre dependent recombinase method, these mice express the fluorescent marker tdTomato under the promoter of the vesicular transporter of GABA.

Animals were kept under controlled temperature of 23 ± 1 °C and photoperiod of reversed 12 h light/dark cycle (light available from 8p.m. to 8a.m.) conditions and group-housed in standard cages with environmental enrichment elements. Food and water were provided *ad libitum*. Females were weaned at 20-21 days of age and group-housed with two to five animals.

After reaching 6 weeks of age, females were exposed to adult C57BL/6 male soiled bedding once per week to stimulate the natural reproductive cycle. All procedures were carried out in accordance with the animal protocols approved by the Portuguese National Authority for Animal Health (*Direcção Geral de Alimentação e Veterinária*; DGAV) and the Commission for Experimentation and Animal Welfare of the Champalimaud Centre for the Unknown (*Órgão para o Bem Estar Animal*; ORBEA).

### Reproductive Cycle Monitoring

To assess the reproductive state of naturally cycling female mice, vaginal cytology samples were collected every morning for at least one entire cycle (four/five consecutive days). This collection was done by vaginal lavage using a pipette (T-210-Y, AXYGEN, with its tip cut): 10μL of 0.01M phosphate-buffered saline (PBS) was gently flushed into the vagina and back out three or four times without touching the vaginal wall to avoid cervical stimulation and pseudopregnancy. The flush containing vaginal fluid was transferred to a glass slide and dried (Caligioni, 2009). Papanicolaou staining was used to differentiate cells (Bio-Optica protocol) and they were observed under a Zeiss AxioScope A1 brightfield microscope with a 10x objective. The identification of the reproductive cycle phase was done based on the proportion of each cell type in the smear. Proestrus is characterized by nucleated epithelial cells. In estrus the main cells present in the vaginal secretion are cornified squamous epithelial cells, clustered with irregular shape. These start to be replaced by leukocytes during metestrus, being these predominant cells in diestrus (Caligioni, 2009; Pfaus, 1999; Snoeren, 2018). Experiments were performed when females were in proestrus/estrus (sexually receptive) or in diestrus (sexually non-receptive).

### *Ex vivo* Electrophysiological Recordings

PR-Cre-EYFP mice were deeply anesthetized with isoflurane. After decapitation, brains were quickly removed and placed into “ice cold” slicing solution containing (in mM): 0.66 kynurenic acid, 3.63 pyruvate, 2.5 KCl, 1.25 NaH2PO4, 26 NaHCO3, 10 D-Glucose, 230 Sucrose, 0.5 CaCl2, 10 MgSO4, and bubbled with 5% CO2 and 95% O2. Coronal sections with 300μm containing the VMH were cut using a vibratome (Leica VT1200) in the same “ice cold” slicing solution. Slices were recovered in oxygenated artificial cerebrospinal fluid (ACSF) containing (in mM): 127 NaCl, 2.5 KCl, 25 NaHCO3, 1.25 NaH2PO4, 25 D-Glucose, 2 CaCl2, and 1 MgCl2, at 34°C for 30 minutes and stored in the same solution at room temperature before being transferred to the recording chamber.

Whole-cell recordings were made under a SliceScope Pro (Scientifica) microscope with the slices submerged in ACSF. Patch recording pipettes (resistance 3-5MΩ) were filled with internal solution containing (in mM): 135 K-Gluconate, 10 HEPES, 10 Na-phosphocreatine, 3 Na-L-ascorbate, 4 MgCl2, 4 Na-ATP, 0.4 Na-GTP (pH 7.2 adjusted with NaOH and osmolarity ~292mOs) and 0.1% of biocytin. Internal solution was filtered with a 0.2μm pore size cellulose acetate filter tip. A Multiclamp 700B amplifier and digitized at 10KHz with a Digidata 1440a digitizer (both from Molecular Devices) were used and the data was filtered online with a 10KHz low-pass filter.

We applied a test-pulse of −10mV for 100ms in voltage-clamp mode at −70mV to monitor membrane capacitance and series resistance. To determine excitability and firing properties, current-clamp experiments were performed at resting potential by applying series of 1000ms current pulses between 0 and 240pA with an increasing interval of 30pA at a frequency of 0.1Hz. Hyperpolarization-activated inward current (Ih current) and rebound firing were measured with a 1000ms current pulse at −90pA. The bridge balance was semi-automatically compensated using Axon pCLAMP 10 (Molecular Devices).

After recording, the pipette was carefully withdrawn from the neurons to allow the resealing of the membrane and the diffusion of biocytin inside the neuron. Slices were then transferred to a multi-well plate containing 4% paraformaldehyde (PFA) for ~1 hour for fixation and then transferred to 0.01M PBS for storage prior to immunostaining.

### Histology

To visualize biocytin filling, slices were incubated in streptavidin-Alexa Fluor 488 Conjugate (Invitrogen; concentration: 1:200) with 0.3% Triton X-100 in 0.01M PBS at room temperature for 3 hours in dark. After washing three times with 0.01M PBS for 30 minutes each, sections were mounted in glass slides (Menzel-Glazer), coverslipped (Marienfeld) with Mowiol mounting medium (Sigma-Aldrich) and the edges sealed with clear nail polish.

For tissue collection, VGat-Cre-tdtomato mice were deeply anesthetized with a lethal amount of a mixture of 12% of ketamine (Imalgene 1000, Merial) and 8% of xylazine (Rompun 2%, Bayer) in saline solution and perfused transcardially with 0.01M PBS followed by 4% PFA in PBS. The brains were removed, fixated in the 4% PFA solution for ~1 hour and transferred to a 30% sucrose (Sigma-Aldrich) in 0.01M Phosphate-Buffer and 0.1% Sodium Azide (ACROS Organics) to allow cryopreservation. Coronal sections with 45μm were cut from the VMH on a sliding microtome (SM2000R, Leica). Sections were imaged with a Zeiss LSM 710 confocal laser scanning microscope with a 10x and a 25x magnification objectives.

### Electrophysiological Data Analysis

A total of 144 recorded neurons were used in the present study. Please see Table 1 and Figure 1 for the numbers of recorded PR+ and PR- neurons, across the reproductive cycle and AP axis.

**Table 1.** Passive membrane properties of the PR+ and PR-populations across the reproductive cycle and the AP axis. Values expressed as Median (IQR).

| *PR+ neurons* |  |  |  |  |  |  |
| --- | --- | --- | --- | --- | --- | --- |
|  | Non-receptive |  |  | Receptive |  |  |
|  | Anterior<br>(n=10) | Medial (n=8) | Posterior<br>(n=11) | Anterior<br>(n=9) | Medial (n=11) | Posterior<br>(n=11) |
| Capacitance<br>(pF) | 9.03 (2.31) | 11.57 (2.66) | 7.34 (5.02) | 11.45 (3.04) | 10.04 (2.256) | 9.57 (2.70) |
| Membrane<br>Resistance<br>(MΩ) | 970.64<br>(1012.68) | 788.77<br>(384.09) | 912.00<br>(1209.69) | 998.98<br>(604.43) | 823.46<br>(853.00) | 1082.56<br>(687.92) |
| Time Constant<br>(ms) | 0.31 (0.09) | 0.36 (0.16) | 0.22 (0.08) | 0.36 (0.18) | 0.27 (0.20) | 0.24 (0.16) |

| *PR- neurons* |  |  |  |  |  |  |
| --- | --- | --- | --- | --- | --- | --- |
|  | Non-receptive |  |  | Receptive |  |  |
|  | Anterior<br>(n=16) | Medial (n=10) | Posterior<br>(n=10) | Anterior<br>(n=11) | Medial (n=11) | Posterior<br>(n=13) |
| Capacitance<br>(pF) | 11.04 (4.70) | 10.05 (1.89) | 8.82 (2.53) | 10.54 (2.90) | 9.54 (1.28) | 10.53 (2.20) |
| Membrane<br>Resistance<br>(MΩ) | 808.55<br>(438.933) | 1025.83<br>(425.53) | 861.21<br>(284.09) | 1258.25<br>(1412.35) | 884.96<br>(654.043) | 871.47<br>(556.16) |
| Time Constant<br>(ms) | 0.36 (0.19) | 0.34 (0.18) | 0.35 (0.17) | 0.36 (0.08) | 0.30 (0.11) | 0.28 (0.16) |

**Figure 1.**
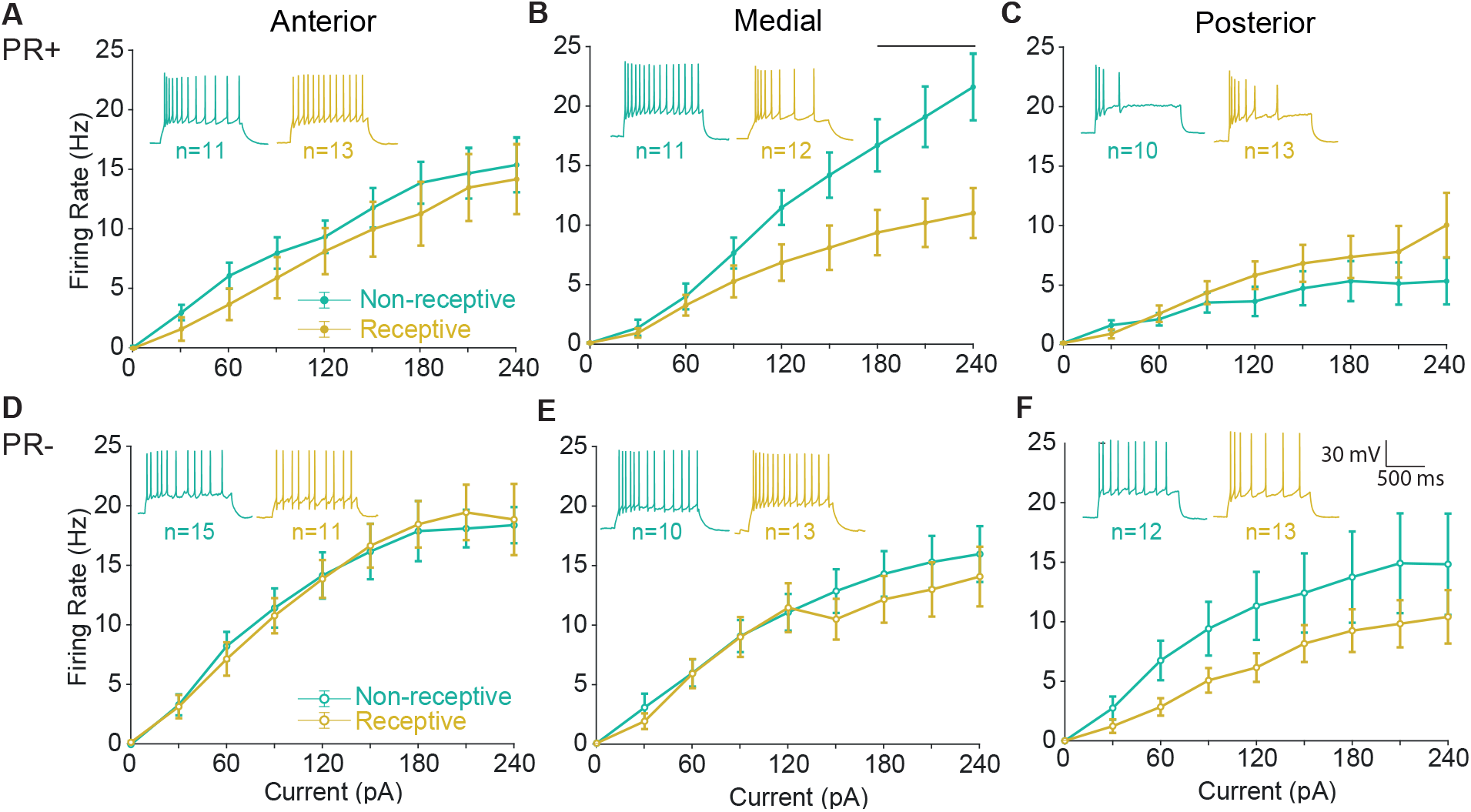
The excitability of medial PR+ neurons changes across the reproductive cycle. (A-C) Input-output (IV) curves across the reproductive cycle reporting the firing frequency of PR+ neurons after injecting 1000ms current pulses between 0 and 240pA (30pA steps), reporting changes only in (B) medial PR+ neurons (p<0.0001, multiple comparison p=0.02 at 180pA, p<0.01 at 210pA, p<0.001 at 240pA). Insets: representative recordings of voltage response from PR+ neurons in response to the same current injection. (D-F) Input-output (IV) curves across the reproductive cycle reporting no changes in the firing frequency of PR- neurons after injecting 1000ms current pulses between 0 and 240pA (30pA steps). Insets: representative recordings of voltage response from PR- neurons in response to the same current injection. (A,C,G,I) PR+ neurons are less excitable than PR-in the anterior and posterior VMHvl (p<0.001 and p<0.01, respectively) but not in (B,H) the medial. Mixed-effects test with repeated measures. Error bars represent SEM.

The action potentials recorded in current-clamp mode were analyzed with custom-written MATLAB code. Each spike within the spike trains obtained upon current injection were detected and used to determine the firing frequency, interspike interval (ISI), threshold to spike, latency to spike and coefficient of variation (CV2= 2|ISI_n+1_ -ISI_n_|/(ISI_n+1_ + ISI_n_)), calculated at a fixed current injection of 150pA) (Holt et al., 1996). The rebound action potentials observed after injecting hyperpolarizing currents were obtained using the same method.

The first action potential obtained in each train was used to calculate the spike amplitude and spike half-width to prevent frequency dependent artifacts in the spike shape caused by sodium and potassium channel inactivation.

The resting potential was determined by averaging the membrane voltage values before the current injection and the Ih current amplitude was calculated by the difference between the early and late phase of a hyperpolarizing current pulse.

Voltage clamp test-pulses were analyzed using Clampfit 10.7 Software (Molecular Devices, LLC) and access resistance (Ra), membrane resistance (Rm), time constant (τ) and capacitance were calculated. Only neurons with access resistance 1/10^th^ lower than the membrane resistance were used for analysis. Access resistance was comparable across location, genotype and the phase of the cycle in the present study.

The passive membrane properties membrane resistance (Rm), membrane capacitance (Cmem), and membrane time constant (τ) were obtained immediately after membrane rupture, using a square voltage step (−10 mV, 100 ms). The access resistance was determined by measuring the amplitude of the current response to the command voltage step and the membrane resistance as the difference between the baseline and the holding current in the steady state after the capacitive decay, by applying the ohm’s law. The membrane time constant was determined by a single exponential fit of the decay phase in response to the square pulse. An approximation of the capacitance was made by using the formula τ = membrane capacitance * access resistance.

### Neuronal Reconstruction and Image Data Analysis

A total of 180 neurons VMHvl neurons were filled with biocytin in acute slices obtained from PR-EYFP female mice using whole-cell recording pipettes. These slices containing biocytin filled neurons were fixed and immunostained for the subsequent reconstruction of neuronal structure. Please refer to Table 2 and Figure 6 for the numbers of filled PR+ and PR- neurons across the reproductive cycle and AP axis.

**Table 2.** Morphological profile of PR+ and PR- neurons across the reproductive cycle and the AP axis. Values expressed as Median (IQR).

| *PR+ neurons* |  |  |  |  |  |  |
| --- | --- | --- | --- | --- | --- | --- |
|  | Non-receptive |  |  | Receptive |  |  |
|  | Anterior<br>(n=18) | Medial (n=14) | Posterior<br>(n=20) | Anterior<br>(n=11) | Medial (n=17) | Posterior<br>(n=14) |
| Soma area ($\mu\text{m}^2$) | 184.88<br>(68.97) | 217.01<br>(95.72) | 193.08<br>(53.31) | 219.77<br>(75.28) | 182.92<br>(38.98) | 219.31<br>(53.94) |
| Dendrites/neuron | 7.00 (2.00) | 7.00 (3.00) | 6.00 (3.50) | 7.00 (3.00) | 9.00 (3.00) | 7.50 (1.00) |
| Primary dendrites/neuron | 4.00 (1.00) | 4.00 (1.00) | 3.50 (1.00) | 4.00 (1.00) | 4.00 (1.00) | 3.50 (1.75) |
| Branch points/neuron | 3.00 (2.00) | 3.00 (3.00) | 3.00 (3.00) | 4.00 (3.00) | 5.00 (4.00) | 4.00 (1.75) |
| Dendritic length of LPDs | 572.69<br>(199.05) | 448.43<br>(268.22) | 337.53<br>(287.65) | 640.69<br>(277.72) | 473.32<br>(273.33) | 482.57<br>(202.10) |
| Average dendritic length of SPDs | 205.36<br>(102.82) | 214.50<br>(65.76) | 145.53<br>(93.24) | 181.54<br>(71.97) | 170.55<br>(77.82) | 177.85<br>(75.57) |
| *PR- neurons* |  |  |  |  |  |  |
|  | Non-receptive |  |  | Receptive |  |  |
|  | Anterior<br>(n=16) | Medial<br>(n=12) | Posterior<br>(n=13) | Anterior<br>(n=10) | Medial<br>(n=10) | Posterior<br>(n=13) |
| Soma area ($\mu\text{m}^2$) | 183.64<br>(58.52) | 208.24<br>(66.82) | 171.12<br>(105.02) | 177.00<br>(124.49) | 188.30<br>(112.71) | 184.28<br>(68.46) |
| Dendrites/neuron | 8.00 (4.00) | 7.50 (3.50) | 7.00 (3.00) | 7.00 (1.50) | 7.00 (2.50) | 7.00 (2.75) |
| Primary dendrites/neuron | 4.00 (1.75) | 3.00 (1.00) | 4.00 (2.00) | 3.50 (2.25) | 3.00 (0.75) | 4.00 (1.25) |
| Branch points/neuron | 4.00 (5.50) | 4.00 (3.50) | 3.00 (2.00) | 3.00 (3.25) | 4.00 (1.00) | 3.00 (3.25) |
| Dendritic length of LPDs | 642.64<br>(235.57) | 644.05<br>(96.20) | 509.16<br>(253.81) | 566.10<br>(352.88) | 494.52<br>(289.21) | 389.96<br>(194.99) |
| Average dendritic length of SPDs | 155.87<br>(62.25) | 255.72<br>(102.05) | 172.95<br>(79.60) | 124.89<br>(60.77) | 150.58<br>(134.84) | 176.90<br>(85.55) |

To determine the Bregma coordinate for each reconstructed neuron and ensure that its location fell within the VMHvl, in a separate experiment we first assessed the extension of the VMHvl, taking advantage of the VGAT-tdTomato. The VMH is organized in a core of glutamatergic neurons (Ziegler et al 2002, Yamamoto et al 2018) and a surrounding shell with higher density of GABAergic neurons (Yamamoto et al 2018). Therefore, we assessed the AP extension of the VMH by counting consecutive slices in which the VMH was detectable, as shown by a decrease of the VGAT-tdTomato signal (Fig. 6A) and multiplying by the thickness of the histological slices. A sharp transition from low to high tdTomato fluorescence was observed both in its anterior limit (near the anterior hypothalamus) as well as with its posterior limit (near the premammilary hypothalamus) (data not shown). The extension obtained matched the extension illustrated by the Paxinos Brain Atlas: Anterior limit= 1.05 to posterior limit= 2.06 from Bregma (that is, the VMHvl spans for ~1 millimeter). For each mouse, we systematically sampled 3 consecutive levels of the VMH in slices of 300 microns of thickness. We thus categorized each sampling level as anterior (Bregma −1.05 to −1.30 mm approx.), medial (Bregma - 1.30 to −1.65 mm approx.), and posterior (Bregma −1.65 to −2.00 mm approx.) levels of the VMHv (Fig. 6B-C). Localization of the reconstructed neurons in the brain was assessed by matching in Adobe Illustrator CS6 (Adobe Systems Incorporated) the confocal images of the neurons with adapted sections from the Paxinos brain atlas (Franklin & Paxinos, 2008). Only the neurons inside the defined boundaries of the VMHvl were considered for quantification. We used Simple Neurite Tracer178 (Fiji/ImageJ software179) to analyze the morphological properties and Sholl profiles of the neurons filled with biocytin. The Cell Counter Fiji/ImageJ plugin was used to manually count spines.

Cut and resealed dendrites (identified as a globular thickening in the extreme of a dendrite) were quantified for every neuron and did not reveal significant differences across groups (Number of cut dendrites/total number of dendrites quantified per group for PR+ neurons: 21/126 in D-anterior, 19/120 in PE-anterior, 20/116 in D-medial, 31/151 in PE-medial, 26/131 in D-posterior and 26/121 in PE-posterior. And for PR- neurons: 26/130 in D-anterior, 16/70 in PE-anterior, 19/95 in D-medial, 15/85 in PE-medial, 26/100 in D-posterior and 17/100 in PE-posterior). When tested with a Fisher’s exact test, neither PR+ nor PR- revealed statistical difference in the proportion of cut dendrites. We thus assume that this unavoidable artifact of our method of neuronal recording and reconstruction did not add any group-specific bias in the structural quantification.

In our hands, as well as in previous studies (Calizo & Flanagan-cato, 2000), the intracellular labeling of VMHvl neurons did not always make their thin axonal process visible, and therefore, the axons identified were not considered for analysis.

Graphs were made using custom made MATLAB code or GraphPad Prism 8 Software.

### Statistical Analyses

Statistical analyses were performed using GraphPad Prism 8 Software. Normality of the residuals was tested with the D’Agostino-Pearson omnibus K2 test. When normally distributed, three-way ANOVA tests were performed to compare groups in different phases of the cycle (D vs PE) vs location in the AP axis (anterior vs medial vs posterior) vs genotype (PR+ and PR- neurons), using the Sidak test to correct for multiple comparisons. In the properties whose residuals did not pass the normality test, a logarithmic or square root transformation was applied and normality was consequently reassessed. As specified in the legend, in the properties that the transformations did not make the residuals become normally distributed, we performed a Kruskal-Wallis test followed by a post-hoc test for multiple comparisons: For Figure 1A-F, a Mixed-effects test with repeated measures, and Figure 7A, B a two-way ANOVA with repeated measures were performed. For Figure 4 and 5B, C a Chi-square test was performed. Whisker plots represent median with interquartile range. Error bars represent mean ± SEM. Significance was noted as *p<0.05.

## Results

### Local changes in the intrinsic excitability and threshold to spike of PR+ neurons across the reproductive cycle

In order to investigate the neuronal excitability of PR+ and PR- neurons, we recorded current to voltage input-output response curves (I-V curves) at resting potential in current-clamp mode in acute slices of naturally cycling PR-Cre x EYFP adult female mice. Briefly, the female reproductive stage was determined by vaginal lavage and slices were obtained from females in the least receptive state (diestrus) and the most receptive state (proestrus/estrus). PR+ neurons were identified by their natural fluorescence under the microscope and PR- by the absence of fluorescence (see Methods for details).

In the anterior and posterior VMHvl, the excitability of PR+ neurons was found not to change across the cycle (Fig. 1A and C). In contrast, medial PR+ neurons of non-receptive females showed significantly higher excitability when compared to the excitability of medial PR+ neurons originating from receptive females, particularly in response to higher input current (Fig. 1B, strong interaction between the phase of the cycle and the amount of current injected (Mixed-effects test, p<0.0001)). The modulation in the excitability of the medial neurons across the reproductive cycle was specific to the PR+ population since no changes were observed across the cycle in the PR- neurons at any of the AP levels (Fig. 1D-F).

In addition, we also observed that the PR+ population was less excitable than the PR- in the anterior and posterior levels of the VMHvl (Mixed-effects test, p<0.001 and p<0.01, respectively) but not in the medial subdivision, probably in part due to the increased excitability of medial PR+ neurons from non-receptive females.

It is worth mentioning that the membrane resistance did not vary across experimental groups (Table 1) and only a small decrease was observed in the capacitance of the anterior and posterior PR+ of non-receptive females (three-way ANOVA, location/phase interaction p=0.04). The membrane time constant (τ) also did not vary across the reproductive cycle, however posterior PR+ neurons exhibited a smaller τ when compared to the anterior and medial PR+ population (three-way ANOVA, location p=0.04).

We observed a small decrease in the capacitance of the anterior and posterior PR+ neurons of non-receptive females (three-way ANOVA, location/phase interaction p=0.04). In turn, while the τ does not vary across the reproductive cycle, posterior PR+ neurons have smaller τ when compared to the anterior and medial PR+ population (three-way ANOVA, location p=0.04).

The resting potential of anterior PR+ neurons of receptive females was lower when compared to non-receptive females (Fig. 2A and B, three-way ANOVA, phase p=0.01, multiple comparison antNon-Rec vs antRec p=0.01). However, this difference was not large enough to produce changes in the threshold to spike (Fig. 2C, Kruskal-Wallis test, p=0.06). The latency to spike of PR+ and PR- neurons did not vary across the cycle (Fig. 2F and H). In addition, we observed a mild, yet significant, lower latency to spike of the posterior neurons that was independent of the phase of the cycle and the genotype (Fig. 2F,H, three-way ANOVA, location p<0.01).

**Figure 2.**
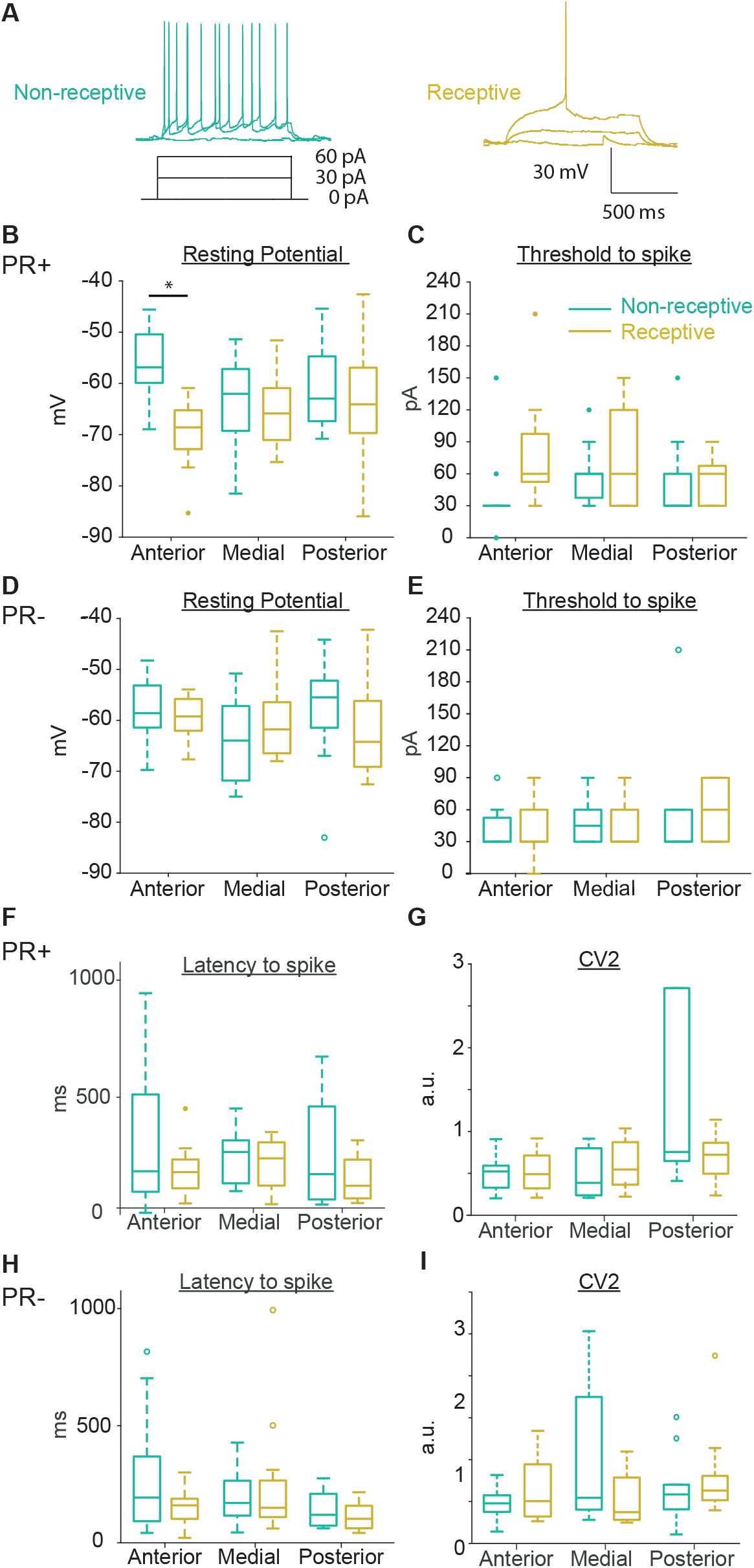
The resting membrane potential of anterior PR+ neurons changes across the reproductive cycle. (A) Representative membrane voltage recordings from PR+ neurons in response to 0, 30 and 60pA. (B) Anterior PR+ neurons of receptive females have a lower resting potential compared to non-receptive females (p<0.01, multiple comparison p<0.01). (C) The threshold to spike, (F) Latency to spike and (G) Regularity of the spike train (CV2) of the PR+ neurons do not change. (D) Resting potential, (E) Threshold to spike, (H) Latency to spike and (I) Regularity of the spike train (CV2) do not vary in PR- neurons across the reproductive cycle.

The regularity of the recorded spike trains was similar across experimental groups, as shown by comparable CV2 values across phase and location for both the PR+ and PR-populations (regularity was determined at 150pA of injected current, Fig. 2G,I).

In focusing on the action potential shape, we observed a comparable amplitude and half-width of the action potentials across the reproductive cycle in PR+ and PR- neurons (Fig. 3A-E). The spike amplitude of the posterior neurons of both PR+ and PR- populations is moderately but consistently smaller (three-way ANOVA, location p<0.01).

**Figure 3.**
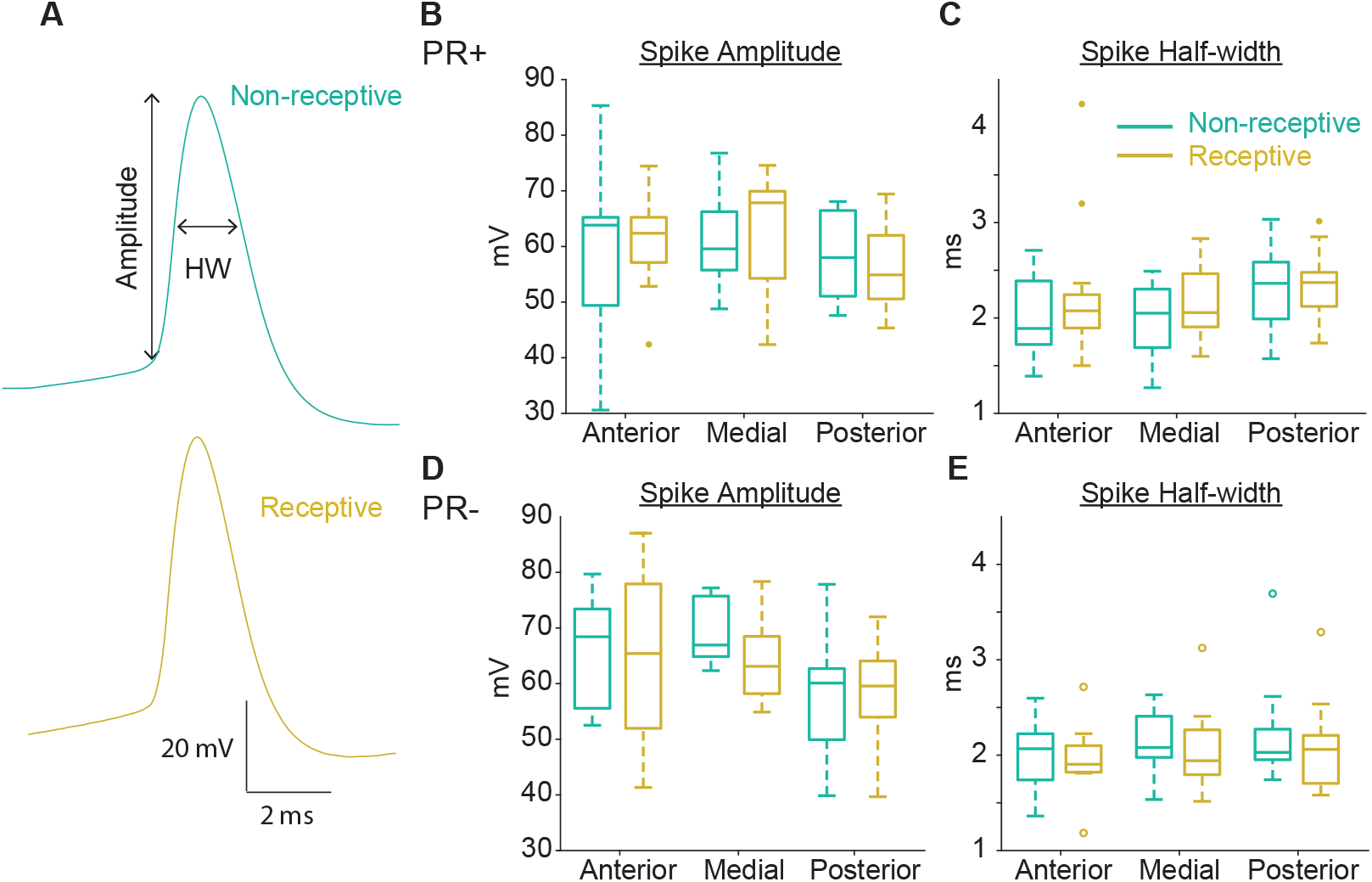
The spike waveform of PR+ and PR- neurons is consistent across the reproductive cycle. (A) Representative examples of the average spike waveform illustrating the amplitude and the half-with (HW). (B) The amplitude of the first action potential for each current injection step of the PR+ neurons does not change across the reproductive cycle nor the AP axis. (D) The spike amplitude of the PR- neurons is maintained across the cycle, however it differs across the AP axis (p<0.01). (C, E) The spike half-width of PR+ and PR- neurons is consistent across the reproductive cycle and the AP axis.

**Figure 4.**
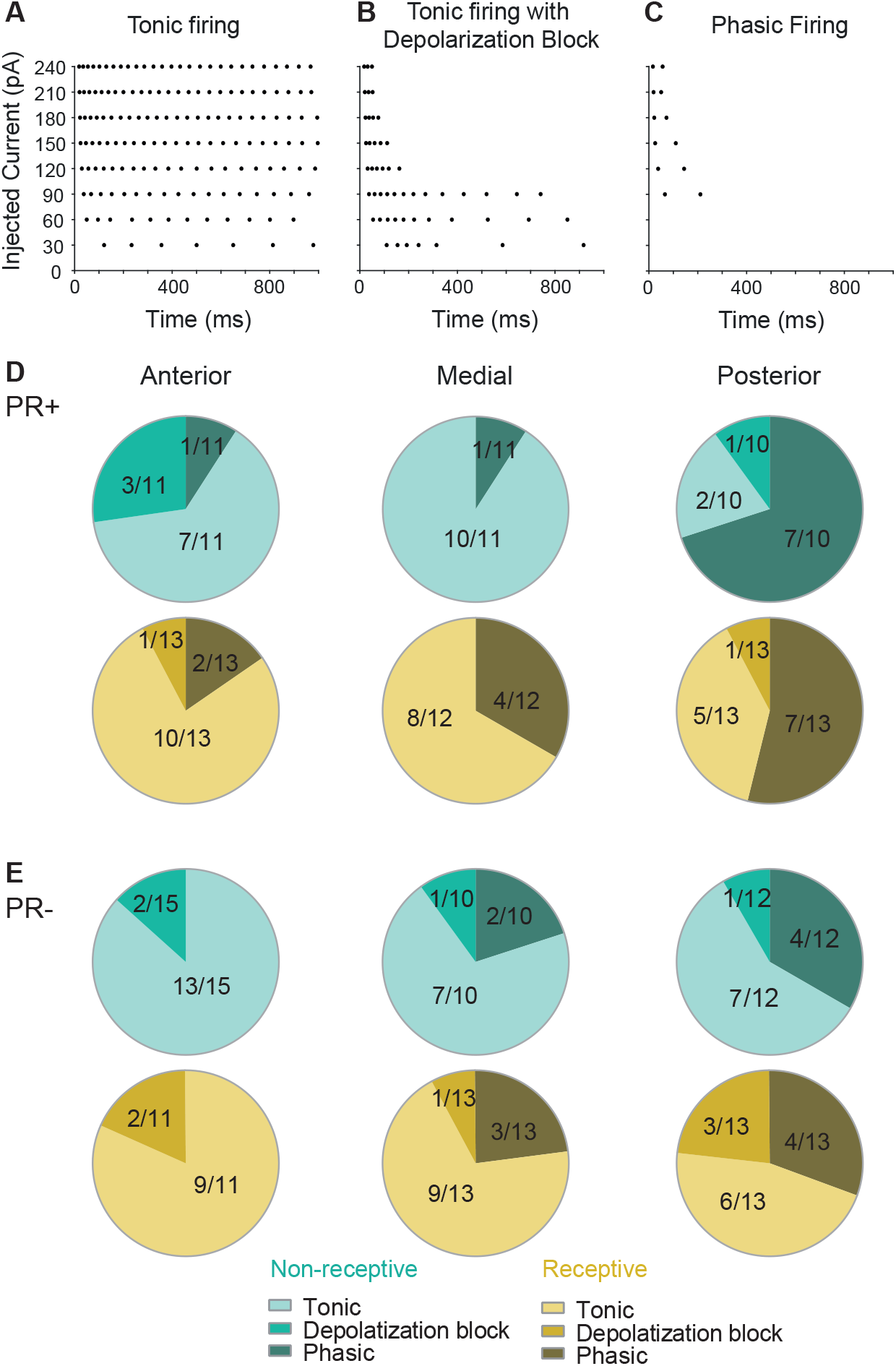
The predominant firing pattern of PR+ neurons varies across the AP axis and differs from the PR-population. Example of (A) Tonic firing, (B) Tonic firing with depolarization block, and (C) Phasic firing patterns in response to injected current steps. (D) The majority of PR+ neurons in the anterior and medial VMHvl display tonic firing, while the posterior PR+ neurons have mainly phasic firing (p<0.001), with no changes across the reproductive cycle. (E) The PR- neurons display mainly tonic firing across the whole AP axis and do not change across the reproductive cycle.

**Figure 5.**
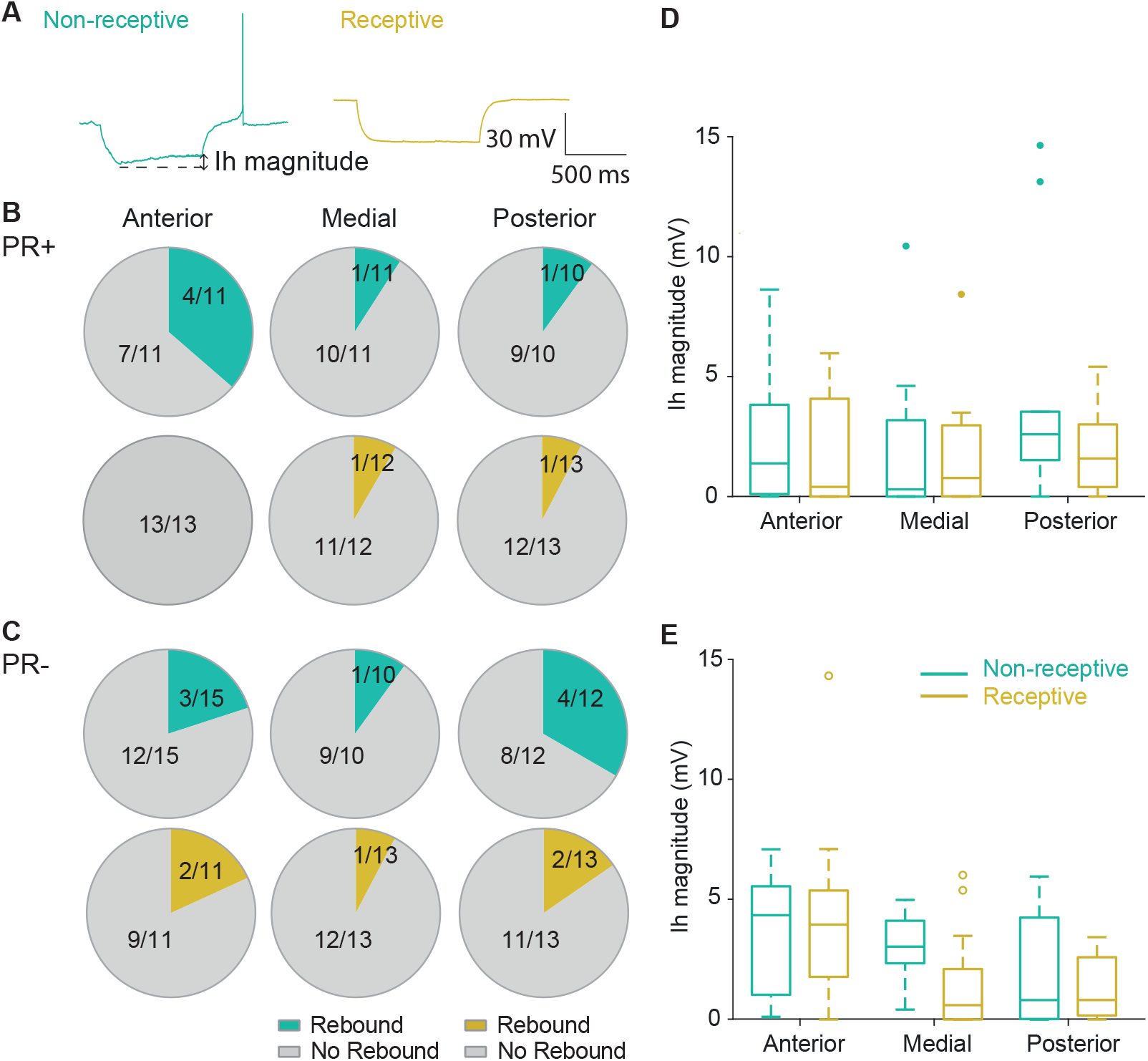
The proportion of anterior PR+ neurons with rebound firing changes across the reproductive cycle. (A) Example of recordings of voltage response at −90pA. Proportion of neurons presenting rebound firing after hyperpolarization of (B) PR+ neurons from non-receptive and receptive females, and from (C) PR- neurons from non-receptive and receptive females. The anterior PR+ neurons of non-receptive females display a significantly higher fraction of neurons with rebound spikes compared to receptive females (p=0.01) (D, E) The Ih amplitude of the PR+ neurons does not change across the reproductive cycle nor the AP axis, while the Ih amplitude of PR- neurons varies across the AP axis (p=0.01).

**Figure 6.**
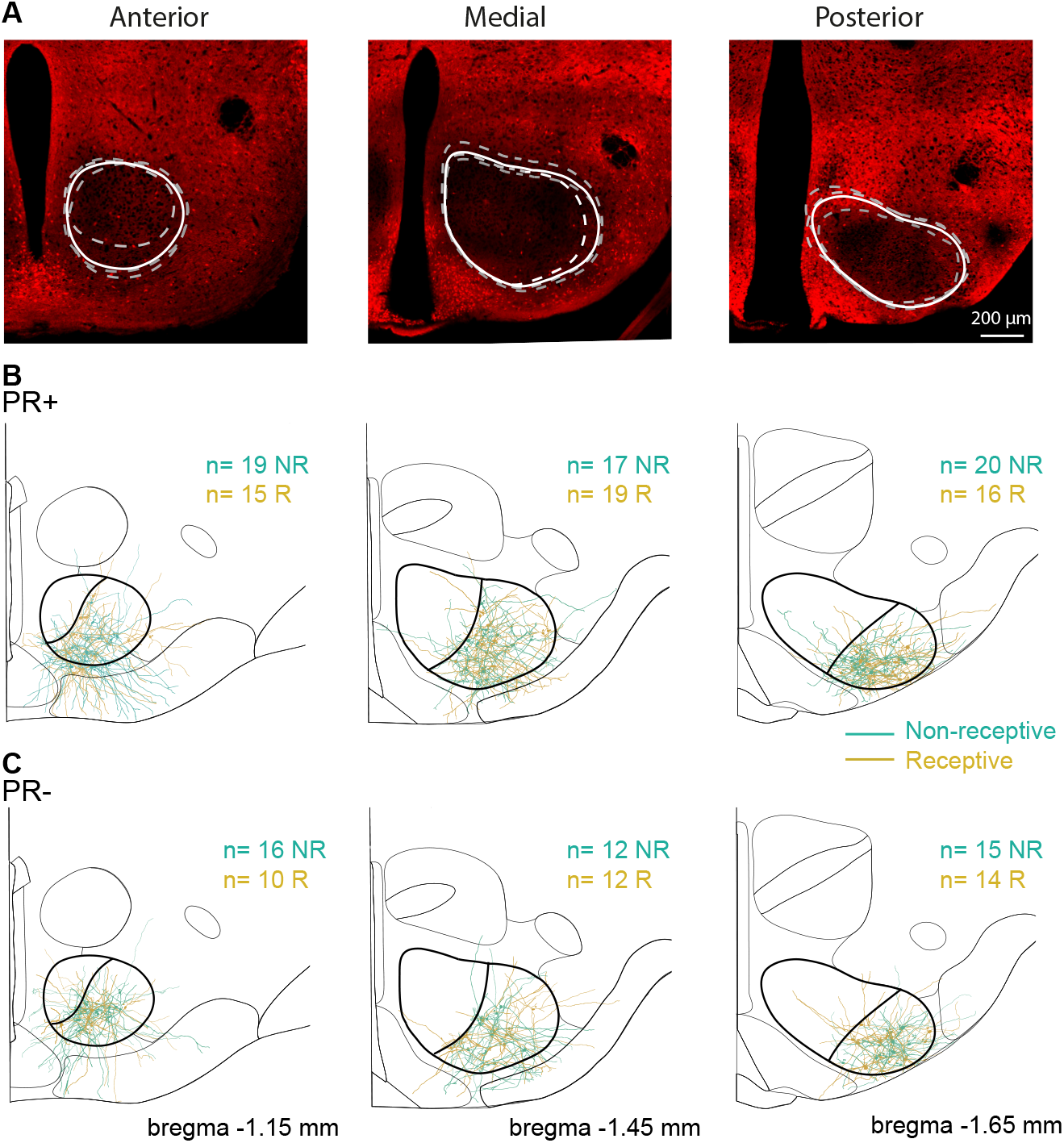
Localization of reconstructed PR+ and PR- neurons across the VMHvl. (A) Limits of the VMH obtained with the inspection of GABAergic markers across the AP axis. Schematic representation of the reconstructed PR+ (B) and PR-(C) neurons and their location in the VMHvl and across the AP axis. Scale bars 200μm.

**Figure 7.**
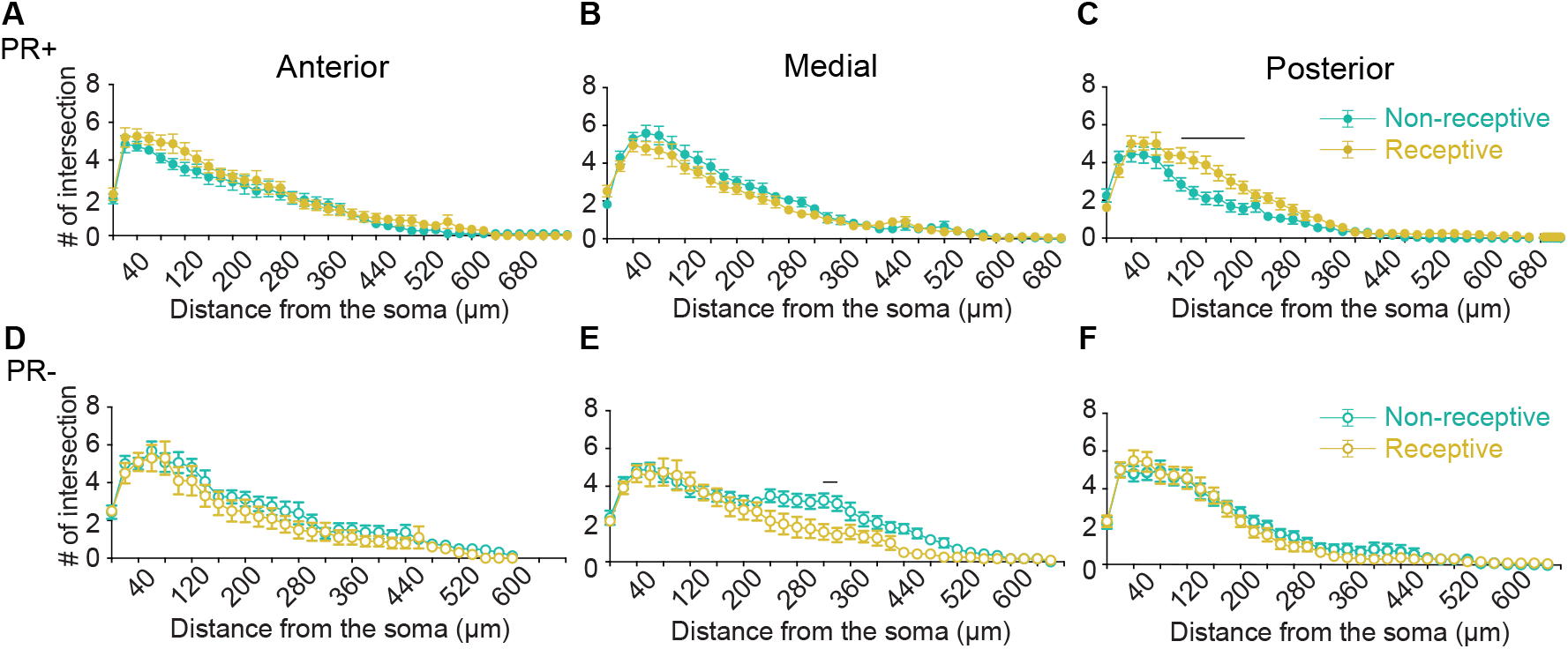
The dendritic properties of PR+ neurons are differently modulated across the reproductive cycle depending on the location in the AP axis and differ from the PR-population. Sholl profiles of (A-C) PR+ and (D-F) PR- neurons across the reproductive cycle. C) The posterior PR+ neurons of receptive females show higher proximal complexity than the ones from non-receptive females (phase of the cycle p=0.01, distance from soma x phase of the cycle p<0.0001, multiple comparison p<0.001 at 120μm, p<0.0001 at 140-160μm, p<0.01 at 180-200μm, and p=0.04 at 220μm. (E) Medial PR- neurons from receptive females show lower complexity than the ones from non-receptive females (distance from soma x phase of the cycle p<0.0001, multiple comparison p=0.03 in340μm). (B, E) The medial PR+ neurons have lower distal branching than the medial PR- neurons (distance from soma x genotype p<0.0001) and (C, F) the posterior PR+ neurons have lower proximal branching than the PR- neurons (distance from soma x genotype p<0.01). Two-way ANOVA with repeated measures. Error bars represent SEM.

Together, these results suggest that the electrophysiological properties of VMHvl neurons vary across the reproductive cycle, but the observed changes depend on their location in the AP axis. In the sexually receptive phase, anterior PR+ neurons are more hyperpolarized in their resting state and the medial PR+ population exhibits lower excitability.

### The proportion of tonic and phasic firing PR+ neurons varies across the antero-posterior axis

In the present study, we observed two major categories of VMHvl neurons according to their firing patterns: tonic and phasic neurons. Tonic firing neurons exhibited a linear increase in firing rate in response to the increases in the magnitude of current injected. Interestingly, within this main class of neurons, some neurons did not reach the depolarization block within the range of currents we injected (Figure 4A, tonic neurons), while some did (Figure 4B, tonic neurons with depolarization block). Amongst phasically firing neurons, action potentials did not linearly increase in response to increased current injection (Figure 4C). These neurons exhibited a step function-like behavior, where after reaching the firing threshold, the number of action potentials remained constant with increased amounts of current. The majority of the anterior and medial PR+ neurons displayed tonic firing, while most of the posterior ones displayed phasic firing (Figure 4D, Chi-square test, p<0.001), with no differences across the reproductive cycle. In contrast, the PR- population is homogenously composed of a higher proportion of tonic firing neurons across the AP axis.

To summarize, posterior PR+ neurons have a higher density of phasic neurons in comparison to the anterior and medial levels of the VMHvl.

### The anterior VMHvl presents a higher proportion of rebound firing neurons during the non-sexually receptive phase of the cycle

Finally, and given that it is well established that hyperpolarizing input can trigger rebound depolarizing responses increasing the firing rate of neurons in a wide variety of brain areas including the hypothalamus (Burdakov et al., 2004; Israel et al., 2008), we sought to characterize the response of VMHvl neurons to hyperpolarizing input. To do so, we quantified the proportion of VMHvl neurons which exhibited rebound firing after hyperpolarization (Figure 5A, −90pA for 1s). We observed that 6 out of 32 PR+ neurons of non-receptive females and only 2 out of 38 PR+ neurons of receptive females fired rebound action potentials (Figure 5B). This difference originates mainly from the anterior PR+ neurons (chi-square test, p=0.01), where 4 out of 11 neurons of non-receptive females displayed rebound firing while in receptive females none of the 13 recorded neurons produced rebound action potentials. Furthermore, no significant differences were observed between PR+ neurons across the AP axis nor between the PR+ and the PR-populations (Figure 5B and 5C).

Rebound depolarization is caused by the hyperpolarization activated inward current (Ih) which represents a powerful modulator of neuronal firing frequency and timing (Momin et al., 2008). Thus, we quantified the Ih current amplitude in VMHvl neurons (Figure 5D and E). Anterior PR- neurons have modestly higher Ih magnitude compared to medial and posterior PR- neurons and to anterior PR+ neurons (three-way ANOVA, location/genotype interaction p=0.01), however, these differences are not reflected in a significantly different fraction of rebound anterior PR- neurons compared to anterior PR+ neurons (Fig. 5B, C).

To summarize, the proportion of neurons with rebound firing is fairly homogenous in the VMHvl across the reproductive cycle and independently of genotype, with the exception of its most anterior subdivision, where PR+ neurons exhibit cyclical alterations.

### Localized structural plasticity of VMHvl PR+ neurons across the reproductive cycle

To study the impact of the phase of the reproductive cycle on the morphological properties of VMHvl neurons, we reconstructed and performed morphometric quantifications of the soma area, number of dendrites, primary dendrites and branch points per neuron of PR+ and PR- neurons in different locations across the AP axis of the VMHvl. As previously reported, the VMHvl is primarily composed of glutamatergic neurons, that are surrounded by GABAergic neurons (Yamamoto et al 2018). In fact, the absence of GABAergic somas has been used to delineate the boundaries of the VMH (Jarvie & Hentges, 2012). Taking advantage of mice expressing the fluorescent reporter tdTomato under the promoter of the vesicular transporter of GABA (VGat-tdTomato) (Kaneko et al., 2018) we observed, as expected, lower signal intensity inside the boundaries of the VMH (Fig. 6A) that was consistent across the AP regions of the VMH. The limits obtained with the inspection of GABAergic markers were used to delineate the boundaries of the VMHvl, and ensure that the neurons characterized in this study were indeed within the nucleus (Fig. 6B and C).

The somatic area was unaltered across the reproductive cycle and was not different between the PR+ and PR- populations (Table 2). We observed that PR+ neurons of receptive females exhibit a robust increase in the number of dendrites per neuron across the AP axis (Table 2, three-way ANOVA, genotype/phase interaction p<0.01), and a higher number of branching points per neuron compared to neurons originating from non-receptive females (Table 2, three-way ANOVA, genotype/phase interaction p=0.01). The changes across the reproductive cycle are specific for the PR+ population, as they were not observed in PR- neurons (Table 2). Both in medial PR+ and PR- neurons, independent of the phase of the cycle, we observed a moderately lower number of primary dendrites per neuron, compared to the anterior and posterior neurons which yielded small yet significant differences across the AP axis (three-way ANOVA, location p=0.04).

Neurons in the VMHvl are characterized by having a long primary dendrite (LPD) that can be several fold longer than its short primary dendrites (SPD) (Calizo & Flanagan-cato, 2002). Therefore, we analyzed LPD and SPD separately (Table 2). While the length of the LPD of PR+ was unchanged across the reproductive cycle, PR- neurons exhibited longer LPDs in non-receptive females (three-way ANOVA genotype/phase interaction p=0.01). No overall significant differences were found in the SPD length of PR+ and PR- neurons across the cycle and AP axis. Interestingly, the posterior neurons were shown to have shorter LPDs than the anterior and medial neurons regardless of the phase of the cycle or the genotype (three-way ANOVA, location p<0.01). No differences in the length of LPDs and SPDs were found between the PR+ and PR- populations.

In addition to the analysis of dendritic length, we sought to investigate whether the complexity of the dendritic trees was modified across the reproductive cycle. To do so, we used the Sholl method that measures the number of dendritic processes intersections as a function of the radial distance from the soma. No changes were observed in the maximum number of dendritic intersections of PR+ and PR- neurons across the phase of the cycle (Figure 7A-F) nor across their location in the VMHvl, indicating that these neurons reach comparable maximum complexities in their dendritic trees. Nevertheless, the analysis of the Sholl profiles of these neurons revealed that the posterior PR+ neurons of non-receptive females had a significantly lower complexity of their dendritic tree (Fig. 7C, repeated measures ANOVA p<0.0001) compared to those from females in the receptive phase. In addition, the medial PR- neurons of non-receptive females had a higher complexity than their counterparts in receptive females (Fig. 7E, repeated measures ANOVA p<0.0001).

The increased complexity observed in the posterior PR+ neurons of females in the receptive phase was large enough to provide differences when tested with the same method specifically in the proximal branching (Fig. 7C, <250μm from the soma), while the increased complexity of the medial PR- neurons of non-receptive females were observed at more distal parts of the soma (240 to 340μm) and were large enough to yield multiple comparisons significant changes (Fig. 7E).

We also observed that medial PR+ neurons have lower dendritic branching than medial PR- neurons (Figure 7B and 7E, repeated measures ANOVA p<0.0001), particularly in the distal parts from the soma (320 to 380μm). The posterior PR+ neurons have lower proximal dendritic branching than the posterior PR- neurons (Figure 7C and 7F, repeated measures ANOVA p<0.01). These observations indicate that PR+ neurons have a different dendritic complexity proximal to the cell body compared to that of the PR-population, independent of the phase of the reproductive cycle.

To summarize, similarly to what we report for the electrophysiological properties, the structural properties of PR+ and PR- neurons are diverse, with some varying across the AP axis and the reproductive cycle.

### Dendritic spine density largely differs between the PR+ and PR- populations

Previous studies have shown that externally primed estrogen exerts effects on the dendritic spine density of VMHvl neurons that are different depending on the type of dendrite, specifically, an increase in spine density on SPDs of VMHvl neurons (Calizo & Flanagan-cato, 2000) and a decrease of the spine density on LPDs of VMHvl neurons expressing the receptor for estrogen (Calizo & Flanagan-cato, 2002). In order to determine if such modulation of the spine density of VMHvl neurons is present in the physiological range of sex hormone fluctuation, the dendritic spine density of PR+ and PR- neurons was analyzed depending not only on their location in the VMHvl, but also on the primary dendrite category across the reproductive cycle of naturally cycling females. Although overall PR+ and PR- neurons have comparable spine densities on SPDs across the reproductive cycle (Fig. 8A and C) and location, posterior VMHvl neurons in non-receptive females have higher spine density on SPDs compared to VMHvl neurons in the receptive phase resulting in a significant interaction between the phase of the cycle and location within the VMHvl (three-way ANOVA, phase/location interaction p=0.02). Furthermore, the PR+ population has significantly lower spine density on the SPDs than PR- neurons (three-way ANOVA, genotype p<0.0001).

**Figure 8.**
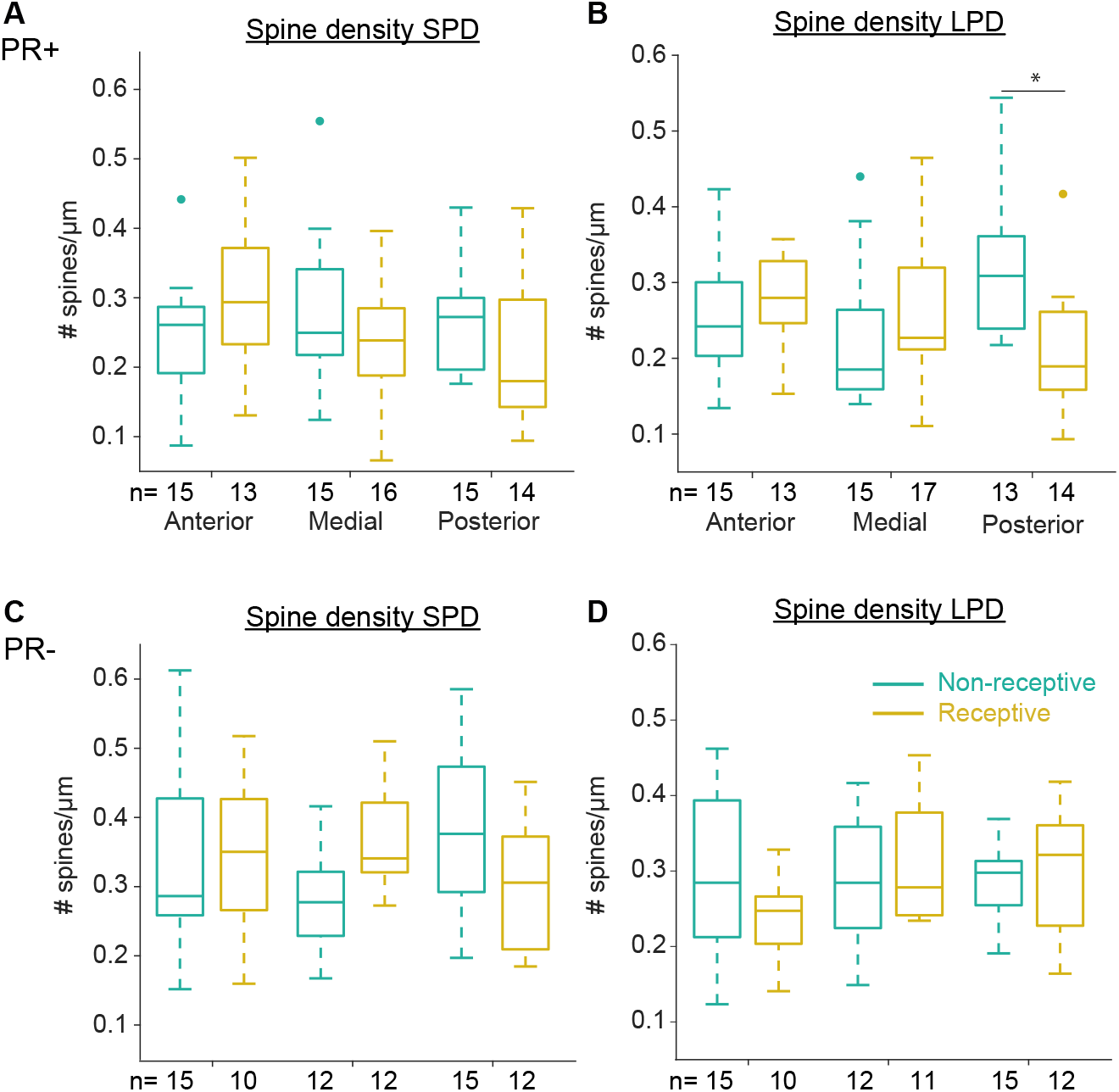
The spine density of the LPD of posterior PR+ neurons changes across the reproductive cycle. (A) Spine density of the SPDs and the (B) LPDs of PR+ neurons across the reproductive cycle. The spine density of the LPDs of posterior PR+ neurons is reduced in receptive females (multiple comparison p=0.0170, causing a change in the interaction between the phase of the reproductive cycle and the location in the AP axis on the LPDs p=0.01). (C) Spine density of the SPDs and the (D) LPDs of PR- neurons across the reproductive cycle. (A,C) PR+ neurons have lower spine density on the SPDs (p<0.0001) and also (B,D) lower spine density on the LPDs (p<0.01) than the PR- neurons. Three-way ANOVA. *p<0.05. Medians ± 25-75 percentile.

Similar to what was observed for the spine densities of SPDs, for LPDs we observed that the posterior VMHvl neurons of receptive females present a decreased spine density compared to posterior neurons from females that were non-receptive, which in the case of LPDs the change is specific to PR+ neurons (Fig. 8B and D, three-way ANOVA, phase/location/genotype interaction p<0.01, multiple comparison posterior PR+ Non-Receptive vs PR+ Receptive, p=0.01).

In addition, by comparing PR+ with PR- neurons, we observed that PR+ neurons have lower spine density on the LPDs in almost all groups, except for the anterior neurons of receptive females and posterior neurons of non-receptive females, in which the opposite is observed (Figure 8B and D, three-way ANOVA, genotype p<0.01).

Altogether, we observed that neurons expressing progesterone receptor have an overall lower spine density when compared to their neighboring PR- neurons. Across the reproductive cycle, posterior PR+ neurons undergo more pronounced structural changes, exhibiting even lower spine density in the receptive phase.

## Discussion

We combined whole-cell recordings with labeling of individual neurons to investigate the structural and electrophysiological properties of PR+ and PR- neurons along the AP axis of the VMHvl and across the reproductive cycle of naturally cycling female mice. We show that PR+ cells are distinct from PR- in three major aspects: first, due to structural properties (PR+ neurons have lower spine densities in general) (Fig. 9B); second, due to the great extent to which the properties of PR+ neurons vary across the AP axis (Fig. 9C); and third due to local changes in the properties of PR+ across the reproductive cycle (which are minimal for the PR- population) (Fig. 9D). Our results further support the existence of subdivisions in the VMHvl and its role in coordinating female behavior with the internal reproductive state.

**Figure 9.**
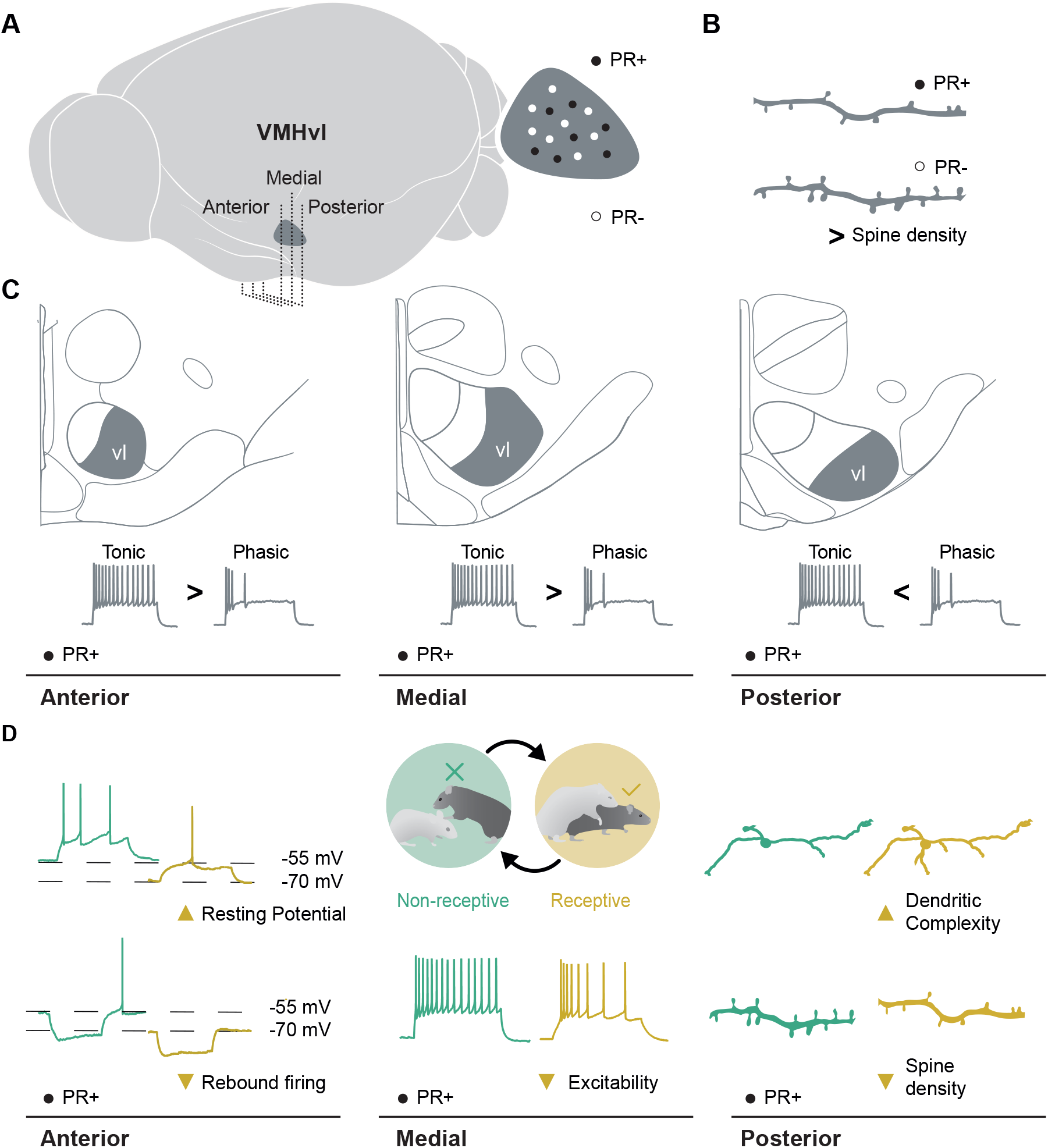
Summary of the structural and physiological properties of genetically defined VMHvl neurons along the AP axis of the VMHvl and its plastic changes across the reproductive cycle. (A) Schematic view of the mouse brain highlighting the location and extension of the VMHvl. (B) The PR+ population was characterized by presenting lower spine density when compared to the PR-population, throughout the whole extension of the VMHvl. (C) The tonic vs. phasic firing proportion of PR+ neurons changes across the AP axis, while it remains constant for PR- neurons. (D) PR+ neurons undergo structural and physiological changes across the reproductive cycle that are specific for the different AP levels of the VMHvl: In the anterior VMHvl, neurons of receptive females have a lower resting potential than the neurons of non-receptive females and do not show rebound firing. Neurons of the medial VMHvl have lower excitability when females are receptive. Posterior neurons undergo structural modulations across the cycle, such as changes in proximal dendritic complexity and spine density.

The impact of the reproductive cycle on VMHvl function has been interrogated *in vivo* and *in vitro*, but most studies were performed in females whose ovaries were removed and then supplemented with estrogen and progesterone (Calizo & Flanagan-cato, 2000, 2002; Griffin et al., 2010; Griffin & Flanagan-Cato, 2008; Millhouse, 1973; Rubin & Barfield, 1980, 1983b, 1983a, but see Inoue et al., 2019; Nomoto & Lima, 2015, for examples of naturally cycling studies). While these manipulations are convenient, easing experimental planning and decreasing interindividual variability (with the extra benefit of allowing a reduction in the number of females used), ovariectomized females with hormonal replacement differ substantially from intact females: first, before hormonal replacement the levels of sex hormones are extremely low, as the main source of estrogen and progesterone is absent and second because the hormonal treatment exposes females to concentrations of sex hormones that differ from physiological levels (Liu & Shi, 2015). The expression of hormone receptors, such as the progesterone receptor, are under the control of sex hormone levels as well (MacLusksy & McEwen, 1978), meaning that that the neuronal response is affected at the level of the receiver (receptor) and message (sex hormone). Persistent hormonal replacement of ovariectomized females leads to tumor development (Kordon et al., 1993) further suggesting that the hormonal treatment leads to undesired physiological effects. Finally, the sexual behavior that hormonally treated females exhibits differs from the behavior of naturally cycling females (Zipse et al., 2000), suggesting that the treatment fails to fully recapitulate the effects of natural sex hormones levels. To the best of our knowledge, this is the first study interrogating the intrinsic properties of neurons across the reproductive cycle of naturally cycling females and therefore the results observed reflect endogenous changes. However, we cannot claim that the changes we observe across the reproductive cycle are an effect of sex hormone levels because those were not directly manipulated. Manipulations with more naturalistic levels of sex hormones or local manipulations in the expression levels of sex hormone receptors should be employed to establish a direct causal link between hormonal levels, the structural and physiological properties of VMHvl neurons and the changes observed.

Our results indicate that PR+ cells are distinct from their neighbors, a difference that is probably established early in development, and later on by the fact that their intrinsic properties of this population can be directly affected by estrogen and progesterone, as PR+ neurons co-express estrogen receptor (Blaustein & Turcotte, 1985; Hashikawa et al., 2017; Sá & Fonseca, 2017). Interestingly, the properties of PR- neurons were similar along the AP axis and across the reproductive cycle, with the exception of changes in the dendritic complexity of neurons in the medial VMHvl. This observation may be explained by the fact that not all ER+ neurons co-express progesterone receptor. ER+/PR- neurons were not labeled with our genetic strategy, but their properties are sensitive to fluctuating levels of estrogen (Calizo & Flanagan-cato, 2000, 2002). Intersectional strategies are needed to address this question.

We observed that the relative proportion of neurons with tonic versus phasic firing changes across the AP axis for PR+ cells, while it remains constant for PR- neurons. This AP gradient may have strong implications not only for the role of different portions of the VMH in controlling behavioral output, but also for the interpretation of activity manipulations, such as the optogenetic control of neural activity. While anterior VMHvl neurons can reach high firing frequencies, posterior phasic neurons present a robust stop of action potential generation after reaching the spiking threshold that does not depend on the current injected in the cell. Importantly, this may be particularly relevant for the interpretation of results where different laser intensities produce diverse behavioral outputs (Kunwar et al., 2015; Lee et al., 2014), as optogenetic activation at particular AP levels may bias the activation of neuronal populations with different intrinsic properties.

Interestingly, even though anterior PR+ neurons had a comparable excitability profile across the reproductive cycle, the resting potential of neurons in the receptive phase was significantly hyperpolarized. This change in resting potential likely underlies a moderate increase in the threshold to fire in these neurons, which required more current to start generating action potentials. Even if this trend did not reach significant differences with our current statistical power, the visual difference between anterior non-receptive and receptive PR+ neurons can hardly be overlooked (Fig. 2C). In addition, the anterior PR+ neurons of receptive females do not show hyperpolarized-induced rebound firing. The fact that these neurons have a more hyperpolarized resting potential, thus more distant from the firing threshold, might explain the fact that the same magnitude of Ih current fails to evoke rebound firing in anterior PR+ at the receptive phase (Fig. 5B). We also report a robust increase of the firing rate under same current injections in medial VMHvl neurons obtained from females in the non-receptive phase compared to those in the receptive phase. These changes were not accompanied by changes in capacitance or membrane resistance across the reproductive cycle of medial neurons, thus it is unlikely that the changes in firing rate are caused by differences in the passive propagation of current into the neuron and instead points towards a modulation of voltage sensitive channels in the medial PR+ neurons, that may be mediated by progesterone (Scharfman & MacLusky, 2006). Overall, these results suggest that the anterior and medial VMHvl have the potential to generate more action potentials in non-receptive females, which is in seeming disagreement with the enhanced-male evoked responses that we previously observed, at the population level, in the VMHvl of sexually receptive female mice (Nomoto & Lima, 2015). However, the results of this study were obtained with extracellular recordings of non-identified neurons and therefore we do not know if they were PR+ or PR-. Also, in the present study, we investigated the intrinsic properties of VMHvl neurons and cannot make any claim regarding the driving input they receive (synaptic and/or neuromodulatory). The VMHvl receives indirect input from the vomeronasal organ, which includes some neurons whose activity is increased in response to male stimuli when females are sexually receptive (Dey et al., 2015). It is conceivable that the enhanced VMHvl activity that we previously observed reflects modifications at the sensory level.

It has been previously reported that hormonal treatment in ovariectomized rats reduced the amount and length of secondary dendrites of a non-defined VMHvl neuronal population (Griffin & Flanagan-cato, 2009). Even if in seeming contradiction with our findings, since we report the opposite effect in dendritic complexity and unchanged dendritic lengths, we would like to point out several possible reasons for such difference. First, the changes we observed are specific to the posterior PR+ population (and absent in our sample of PR- neurons), therefore, sampling VMHvl neurons independently of their genotype could have masked a specific plasticity process in posterior PR+ neurons. Second, in our dataset we observed that medial PR- neurons undergo a reduction in dendritic complexity that goes in line with previous reports. Thus, pooling samples from different levels of the VMHvl may mask changes, or alternatively sampling exclusively in a single AP region of the VMHvl could lead to findings that are limited to that level alone, and thus not generalizable to the whole VMHvl. Interestingly, posterior PR+ neurons did not only undergo changes at the level of their dendritic complexity, but also at the level of spine density, which was reduced in the receptive phase specifically in their LPDs. The fact that such dendritic spine changes are specific to LPDs may suggest that some pathway specific plasticity is happening during the reproductive cycle. Even though it has been previously hypothesized that SPDs may preferentially integrate local excitatory inputs, while LPDs may preferentially interact with long range inputs, inhibitory neurons of the shell and neuropeptides (Griffin & Flanagan-cato, 2011; Yamamoto et al., 2018), as of yet, no circuit mapping study has quantitatively explored what is the anatomical origin of the inputs. Thus, both the anatomical origin of those inputs and their possible relevance for the dendritic coding presented here needs to be further investigated. Our findings suggest a modification of the overall synaptic weights that the LPDs and SPDs will have in the dendritic integration for output generation, with a bias towards the inputs contacting SPDs.

It is worth noting that while the spine densities we observed are comparable to the ones reported in previous studies (Calizo & Flanagan-cato, 2002), the dendritic lengths we report both for PR+ and for PR- neurons are several fold longer than those previously reported (Calizo & Flanagan-cato, 2002). These changes may be explained by the differences in the technical approaches used to investigate neuronal structure. While previous reports have used fixed histological slices (100-150 microns) filled *a posteriori* with lucifer yellow, in the present study we have used acute slices for *ex vivo* electrophysiological recordings (300 microns), in which neurons were filled with biocytin during their electrophysiological monitoring. Thus, several key differences may explain the contradictory results: first, the thickness of the slice, which could allow us to track dendrites for longer extensions; second, the fact that neurons in acute slices for *ex vivo* recordings reseal their membranes, producing a visible thickening in the extremity of their cut dendrites, thus allowing us to recognize incomplete dendrites with our technical approach.

In summary, we have observed a surprising diversity of structural and physiological plasticity processes both along the AP axis of the VMHvl, and across the reproductive cycle, within the genetically defined population of PR+ cells. These findings highlight the repertoire of local plasticity rules across the VMHvl that is probably explained by the different specialized transcriptomic clusters previously characterized across the AP extension of the VMHvl (Kim et al., 2019; McClellan et al., 2006), supporting the existence of subdivisions in the VMHvl and a more complex role of this hypothalamic structure in socio-sexual behavior, which has been recently acknowledged (Hashikawa et al., 2017; Inoue et al., 2019; Sakurai et al., 2016; Wang et al., 2019; Yang et al., 2013). The existence of multiple subdivisions that undergo local changes in response to sex hormone levels hint to the hypothesis that this hypothalamic structure might be able to control a complex and coordinated set of responses in reaction to an ubiquitous message, the systemic levels of sex hormones in circulation, from metabolism to sexual receptivity. This idea is supported by the connectivity of ER+ neurons, which present a high degree of output divergence, with anterior ER+ neurons projecting to pre-motor areas, while posterior ER+ are engaged in amygdalo-hypothalamic loops (Lo et al., 2019). Similar circuit motifs have been described in other hypothalamic nuclei, such as the organum vasculosum of the lamina terminalis (OVLT), where different subnuclei can coordinate a multipronged response in reaction to thirst (Graebner et al., 2015). Altogether, it becomes evident that acknowledging the different neuronal properties within the AP axis of the VMHvl is crucial to have a holistic understanding of its involvement in behavior and further studies with focal manipulations are needed to understand the functional role of observed local properties and their modulation by the reproductive cycle.

## Conflict of interest statement

The authors declare no competing interests.

## Acknowledgments

We thank the Lima Laboratory and Champalimaud Research members for helpful comments on the manuscript, and Gil Costa for the summary figure design. This work was supported by the Champalimaud Foundation, UIDB/04443/2020, LISBOA-01-0145-FEDER-022170, LISBOA-01-0145-FEDER-022231 and an ERC Consolidator Grant (772827, SQL).

## Author contribution

ICD, NG and SQL designed the research. ICD, NG and LF performed the research. ICD and NG analyzed the data. ICD, NG and SQL wrote the paper.

## Notes

### Competing Interest Statement

The authors have declared no competing interest.

